# GW182 Proteins Restrict Extracellular Vesicle Mediated Selective Export of Used miRNAs In Mammalian Cancer Cells

**DOI:** 10.1101/2020.09.11.294488

**Authors:** Souvik Ghosh, Yogaditya Chakrabarty, Kamalika Mukherjee, Bartika Ghoshal, Suvendra N. Bhattacharyya

## Abstract

MicroRNAs are small regulatory RNAs of relatively long half-life in non-proliferative human cells. However, in cancer cells the half-lives of miRNAs are comparatively short. To understand the mechanism of rapid miRNA turn over in cancer cells, we explored the effect of “usage” of specific miRNAs for translation repression of their targets on the abundance of that miRNA. We have noted an accelerated extracellular vesicle mediated export of “used” miRNAs in mammalian cells and the miRNA-export process get retarded by Ago2 interacting protein GW182B. The GW182 group of proteins are localized to GW182 Bodies or RNA Processing Bodies in mammalian cells and GW182B dependent retardation of miRNA export depends on GW-body integrity and is independent of HuR mediated auxiliary pathway of miRNA export. Our data thus support existence of a HuR independent pathway of miRNA export in human cells that can be targeted to increase cellular miRNA levels in cancer cells to induce senescence.

**Graphical Abstract:** 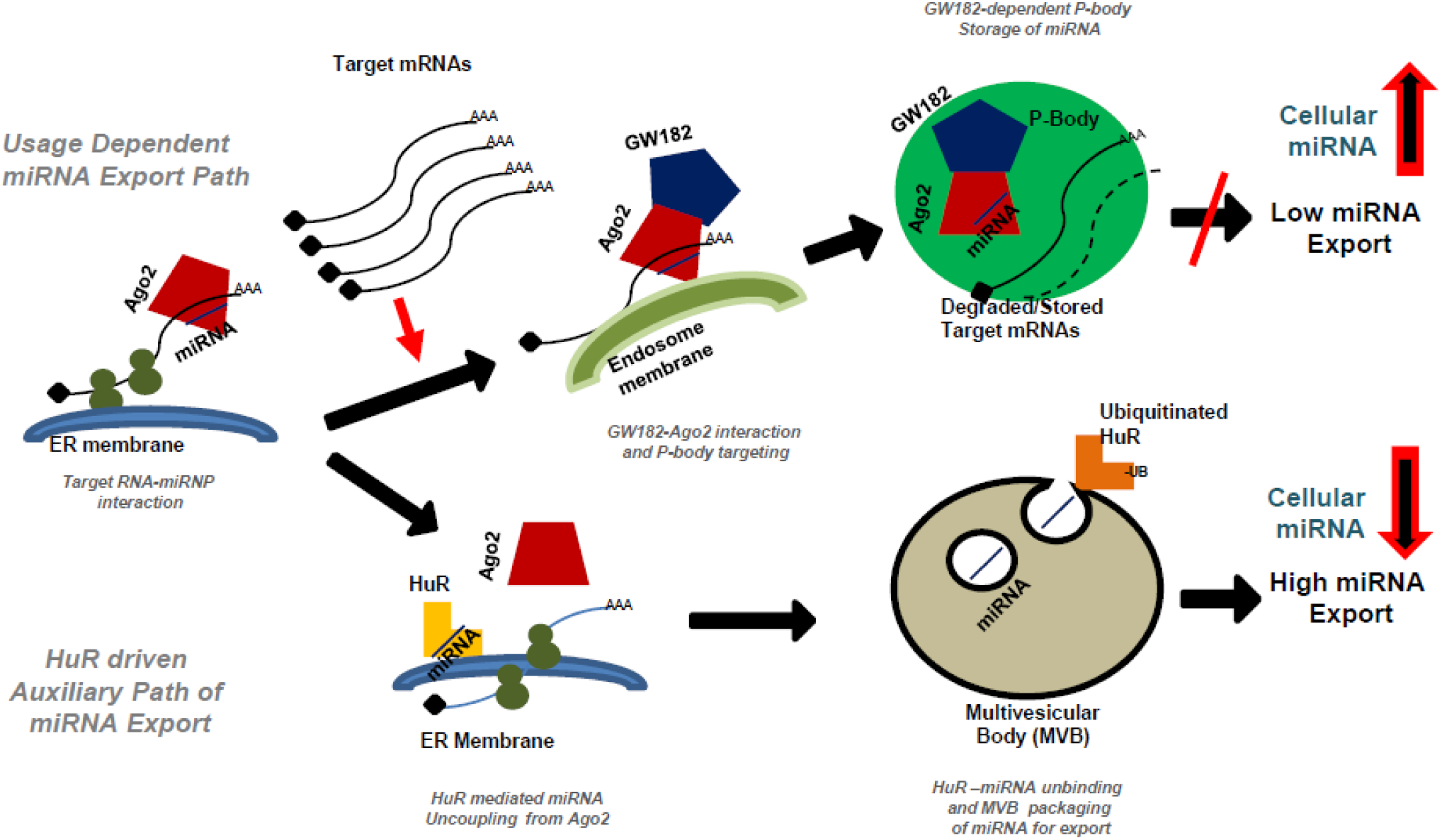

**Key Points:**

I. Usage of miRNA is proportional with its export and turnover
II. GW182 proteins cause phase separation of Ago2 to P-bodies and restrict miRNA export via extracellular vesicles (EVs)
III. HuR driven auxiliary path of miRNA export is independent of GW182 controlled main path of miRNA exclusion

## Introduction

miRNAs, the 22 nucleotide (nt) long non-coding RNAs, forms complex with Argonaute group of proteins and regulate majority of genes by imperfect base pairing to the 3’UTR of target messages [1]. Major fraction of human genes is regulated by one or more miRNAs and this regulation is contextual. Impairment of miRNA function has been documented in human diseases and abnormal expression of miRNAs have been associated with diseases including different types of cancers [2–4].

miRNPs repress target genes by inhibiting protein translation and affecting the target mRNAs stability [5, 6]. Expression and activities of miRNAs have been noted to be sensitive to cellular context [7, 8]. But despite being the important regulator of gene expression in eukaryotes, factors controlling miRNA stability and turnover in mammalian cells are only partly explored.

Mechanism to regulate miRNA level and activity, for the most part, remains limited to discoveries related to specific miRNAs [9]. Recent reports suggest that the RNAs with miRNA binding sites not only influences its activity by altering stability of the corresponding miRNAs in animal cells but also affect its de novo biogenesis [10]. The claim of miRNA stabilization due to its base pairing with target-mRNA in *C. elegans* [11] is opposite to what has been reported in mammalian cells where cellular and viral target messages can downregulate corresponding miRNAs [12–14]. TDMD or Target RNA-Directed MicroRNA Degradation is another pathway described in recent time in mammalian neuronal cells where microRNA 3′-end tailing by the addition of A/U non-templated nucleotides, trimming or shortening from the 3′ end, and highly specific microRNA loss happens [15].

In animal cells, multivesicular bodies (MVB) play an important role in restricting miRNP recycling. Inhibition of MBV formation decreases miRISC activity whereas blockage of MVB maturation into lysosomes leads to increase in miRNP activity [16, 17]. MVBs can also fuse with cell membranes to form exosomes in animal cells [18]. Exosomes, 30-100 nm small extracellular vesicles secreted by the animal cells to the extracellular space are considered as means of intercellular communication [19]. miRNPs are shown to be present in exosomes that serve as intercellular carriers of miRNAs [20].

Extracellular export of excess miRNAs is another way of regulating miRNA levels in human cells. Several post-transcritional modifications of miRNAs are also noted to be linked with their extracellular vesicles (EV)-mediated export. Several RNA binding proteins have been identified as miRNA exporter as they promote export of miRNAs via exosomes or Extracellular vesicle (EV) [21, 22]. Human ELAV protein HuR was identified earlier as a derepressor of miRNA-activity and inhibits the action of miRNA on *cis*-bound target mRNAs primarily by uncoupling the miRNPs from the target messages and mobilizing them from P-bodies in amino acid starved human hepatoma cells [23]. HuR also acts as miRNA sponge and by its reversible binding with miRNAs in stressed cells, it can accelerate miRNA loss by promoting its extracellular export [21].

Phase separation of proteins and other components contribute in regulation of biochemical pathways in mammalian cells and miRNA-regulation has also been found to be compartmentalized in mammalian cells [24]. The memebrane of endoplasmic reticulam (ER) and P-bodies are the compartments where the repression and storage of miRNA targets happen respectively [23, 24]. The process of miRNA or target RNA compartmentalization thus could be the primary mechanism of reversible translation repression and miRNA turnover regulation in mammalian cells.

In mammalian cells, repressed messages can be stored in P-bodies, as the phase separated compartments having increased abundance of miRNA and mRNA catabolising enzymes but where mRNAs can be stored in a reversible manner [23]. GW182 also known as TNRC6 family of proteins has three members in human cells that interact with Ago proteins [25]. GW182 proteins are components of mammalian P-bodies and are the effectors of miRNP mediated gene repression. They also can act as stabilizers of miRNPs [26]. GW182 proteins by recruiting PABP protein prevent the initiation step of protein translation from the target mRNAs [27].

This work shows how phase separation of Ago2, by interacting group of proteins GW182, ensures retardation of miRNA export in mammalian cells. The process depends on the integrity of the separated phase and it opposes the miRNA export that happen otherwise specifically for miRNAs already used for target RNA repression. We observed that the levels of GW182 do not have an effect on HuR driven export of miRNA and thus it acts as the main pathway for miRNA export in human cancer cells other than the known HuR-mediated auxiliary miRNA export path.

## Results

### Usage dependent lowering of miRNAs in human cells

To find out how the usage is linked to miRNA turnover, we expressed reporter mRNAs with three imperfect miRNA binding sites for let-7a and expressed it in HeLa or MDA-MB-231 cells where let-7a miRNA is also expressed. The effects of expression of reporter mRNAs on existing let-7a pools of the respective cells were compared. Quantitative estimation of the remaining let-7a after 48h of reporter mRNA expression was performed. A decrease in existing let-7a level was noted in presence of the target. Similar decrease of miR-122 in hepatic cell Huh7 was also documented in presence of miR-122 target reporter mRNA expressed there (Figure 1A). Similar observation was also documented in HeLa cells when an inducible Renilla luciferase reporter with three let-7a binding sites was expressed there. The reporters were expressed in an inducible manner and after 24h of induction with Doxycycline cellular levels of miRNAs were quantified. Similar observation was there with an inducible miR-122 reporter in Huh7 cells expressing miR-122. In both cases, the non-target miRNAs miR-16 and miR-24 levels were not changed with respective let-7a and miR-122 reporter mRNA induction (Figure 1B). The reduction of the mature miRNA content also dependent on the amount of target mRNA present (Figure 1C). As expected, substrate dependent lowering of a miRNA resulted in higher expression of other endogenous or reporter mRNAs targets of the cognate miRNA in respective cases confirming lowering of effective miRNP pools (Figure S1).

**Figure 1.**
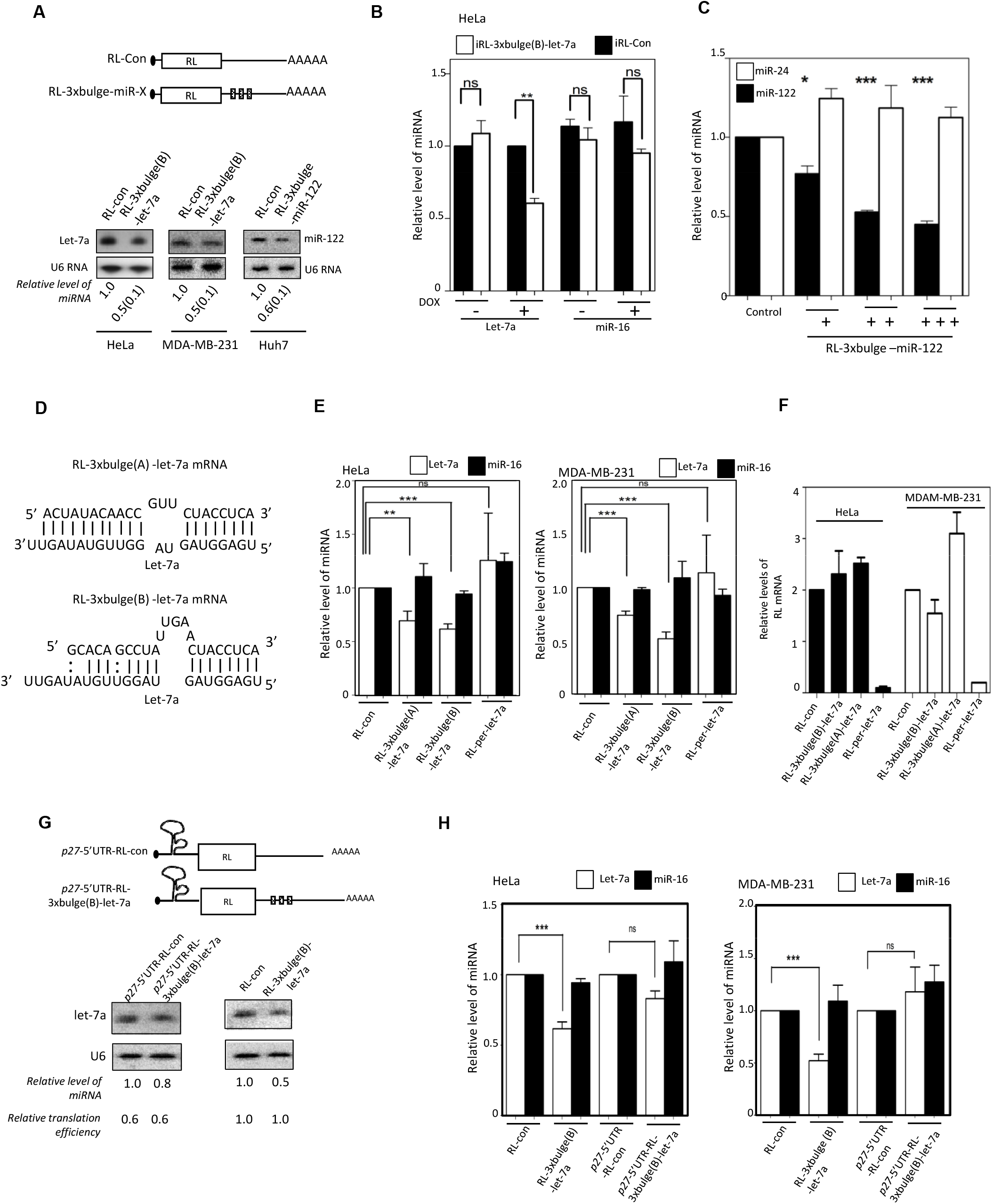
Usage of miRNA reduces its level in human cells. (A) Downregulation in cellular miRNA levels in presence of their substrate mRNAs. Scheme of reporters used for expression of a target message in excess. Representative blots showing let-7a or miR-122 and U6 snRNA levels in human cell lines expressing RL reporters. Quantifications of miRNA levels from multiple experiments (mean+/− s.e.m., n=3 to 5). (B) Relative levels of let-7a and miR-16 in HeLa cells expressing a control or let-7a reporter from the tetracycline inducible constructs. Real time quantifications of endogenous miRNA were done after 24h of induction and were normalized against U6 snRNA. Value obtained with RL-con was set as unit (mean+/− s.e.m., n=3). (C) level of miR-122 and miR-24 in Huh7 cells transfected with increasing concentration (100ng-1000ng for 10^6^ cells) of in vitro transcribed RL reporter mRNA with three miR-122 binding sites. Levels of miRNAs in RL-con (1000ng) (mean+/− s.e.m., n=3). (D) Target site complementarities for two let-7a reporters used to downregulate cellular let-7a level. (E) Level of let-7a and miR-16 in HeLa or MDAM-MB-231 cells expressing control or reporter mRNAs with let-7a binding sites. RL-per-let-7a has one perfect let-7a site. miRNA level normalized against U6 snRNA and values obtained with RL-con reporter were set as 1 (mean+/− s.e.m., n=3). (F) Relative level of RL reporter mRNA levels in HeLa and MDAM-MB-231 cells estimated by real time quantification and values were normalized against GAPDH mRNA level. Values for RL-con set as unit. All experiments were performed in triplicates and values are mean+/− s.e.m. (G-H) Target mRNA containing a translation inhibitory element from 5’UTR of p27 mRNA reduces substrate dependent lowering of let-7a miRNA. Schematic representation of RL reporter construct containing the p27-5’UTR element (G, upper panel). Level of let-7a was measured in HeLa cells expressing either p27- RL-control or p27-RL3xbulge(B)-let-7a mRNAs (G, bottom panel). Luciferase assays for reporter gene and real time quantification of reporter mRNA levels were performed to calculate the relative translation efficiencies defined by amount of protein per unit of mRNA present in reporter mRNA expressing cells. Similar experiments were also done with RL-3xbulge(B)-let-7a and RL-con reporters. Let-7a levels in RL-con reporter transfected cells were considered as 1. Northern blot data in panel G is substantiated with more quantitative data measured by real-time based quantification of let-7a in HeLa and MDAM-MB-231 cells and shown in panel H (mean+/− s.e.m., n=3). ns: non-significant, * P<0.05, **P<0.01, ***P<0.0001. P values were determined by paired t test.

The miRNAs binding to the target sites could be strong or weak depending on the complementarities of target mRNAs to the respective miRNAs. We have used three let-7a reporters; one with one perfect let-7a site (RL-per), the other two have either A or B type bulges and have different complementary to let-7a in non-seed region. The RL-per reporter should undergo a RISC mediated cleavage upon expression in cells expressing let-7a. The A or B type bulge containing reporters are known to get repressed in HeLa cells but known to undergo different levels degradation of the target mRNA [28]. We found the target RNA dependent miRNA decrease has been specific and dependent on target mRNAs availability and on nature of the target sites (Figure 1D-F). The target mRNA with perfect miRNA sites undergoes the RNAi like degradation and availability of target mRNA become less. This target mRNA thus has failed to induce lowering of miRNA content although the respective miRNPs were in use for RISC cleavage (Figure 1D-F). It suggests that usage dependent miRNA lowering very much dependent on active translation of the target messages and therefore the process should also dependent on translatability of the target mRNAs. We found in subsequent experiments that mRNA with a translation inhibitory secondary structure at its 5’UTR [29] caused a reduced downregulation of corresponding miRNAs compared to the same target site containing reporter mRNA without the translation inhibitory sequence in its 5’UTR. This has been tested in two human cell lines (Figure 1G-H) where the let-7a reporter mRNA with p27 5’UTR failed to show downregulation of let-7a. These reporter also showed reduced translatability compared to respective mRNAs without p27 5’UTR.

### Enhanced extracellular export of used miRNA in mammalian cells account for the lowering of miRNA content in human cancer cells

This has been observed in neuronal cells that the lowering of miRNA in presence of its substrate has been resulted from an enhanced degradation of cognate miRNAs due to adenylation of the miRNA that triggers the miRNA degradation by specific exonucleases present in neuronal cells [15]. In human cancer cells, HeLa and MDA-MB-231, we did not documented change in length of the miRNAs in presence of its target (Figure 1A). On contrary, has been reported that extracellular miRNA export could be a key mechanism of miRNA lowering in mammalian cells [30].

Usage related reduction of miRNA could be due to a selective export of “used” miRNAs in human cells. We have noted the extracellular content of miRNA let-7a get increased specifically both in HeLa and MDA-MB-231 cells in presence of abundant amount of translatable target mRNAs. The non-specific miRNA miR-16 and miR-24 did not showed any increase in presence of let-7a target mRNAs in HeLA or MDA-MB-231 cells. The amount of miRNA exported out from HeLa and MDA-MB-231 cells is also dependent on availability of the target messages (Figure 2A). In this context, ectopic expression of a hepatic miRNA in non-hepatic cells, we documented substrate specific export of miR-122 from HeLa cells when its target mRNA was expressed (Figure 2B). We have explored the effect of GW4869, a neutral sphingomyelinase inhibitor that is known for its inhibitory effect on EV-mediated miRNA export [31, 32]. GW4869 blocks EV-mediated export of miRNA from mammalian cells by preventing biogenesis of EVs. The effect of GW4869 on used miRNA export was evident as the lowering of miRNA in presence of its target mRNAs was retarded when GW4869 was applied (Figure 2C-D). SMPD2 is the target of GW48569. Confirming the role of EVs in target dependent miRNA lowering, similar effect was observed on cellular-miRNA content when siRNA-mediated knockdown of the protein SMPD2 was targeted (Figure 2D).

**Figure 2.**
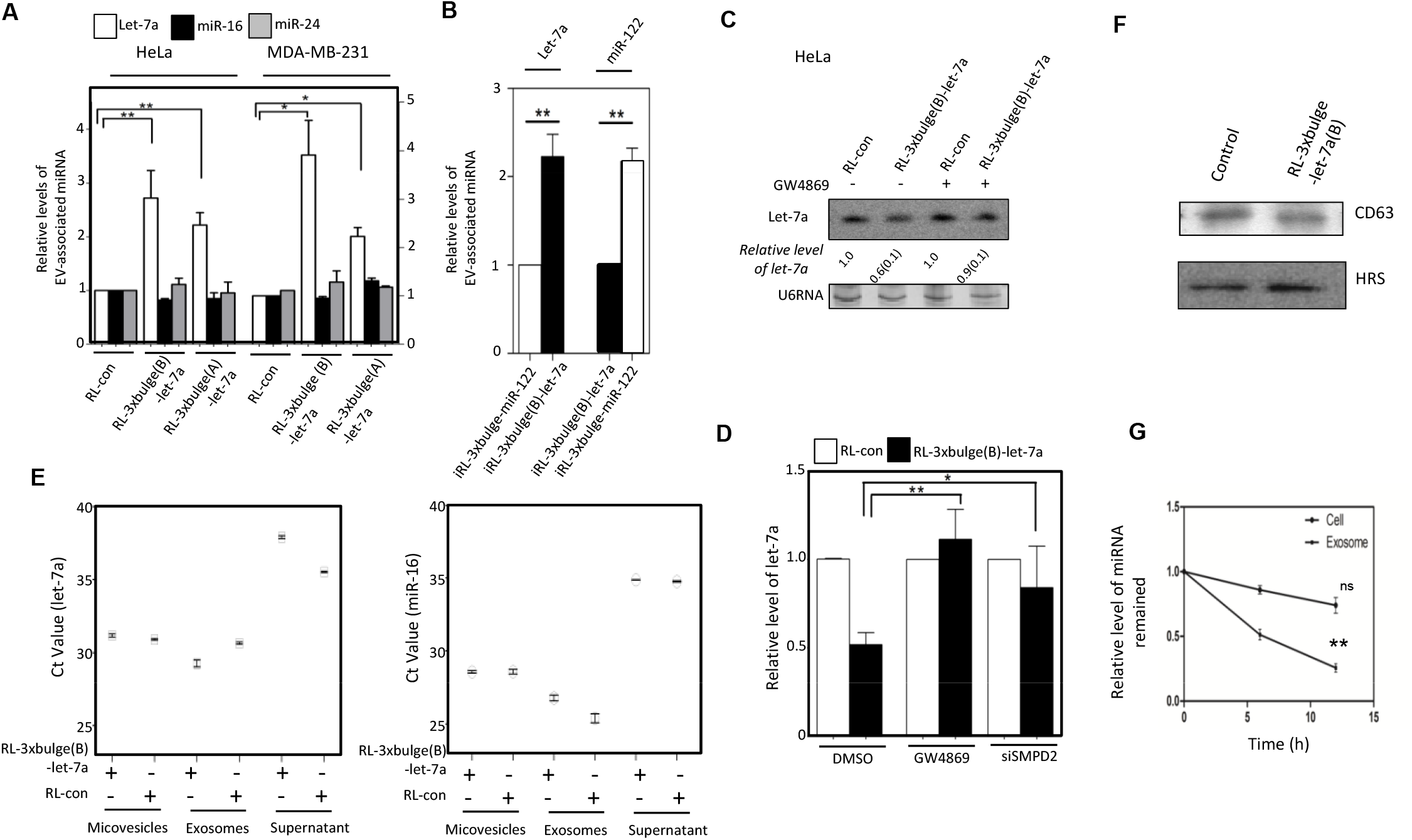
Substrate driven extracellular export regulates cellular miRNA level in mammalian cells. (A) Real time estimations of miRNA levels in EVs isolated from culture supernatants of cells expressing reporter mRNAs with or without let-7a miRNA binding sites (mean +/− s.e.m., n=3). Two different reporters with difference in let-7a complementarities (see Figure 1D) were used. (B) RL control and reporter mRNAs were induced in HeLa cells and cellular level of miR-122 were measured in each case after 24h of induction (mean +/− s.e.m., n=3) (C) Effect of GW4869 treatment on target mediated downregulation of let-7a in HeLa cells. U6 RNA serves as loading control. Quantification was done from three different Northern Blot data. SD value is shown within the bucket. (D) Real time estimation of let-7a levels in MDA-MB-231 cells upon depletion of SMPD2 by RNAi or treatment of control siRNA transfected cell either with DMSO or GW4869 (mean+/− s.e.m., n=4). (E) Relative level of miR-122 present in different extracellular components like microvesicles, exosomes and also in exosome removed culture supernatant. For the same amount of mRNAs, Ct values were plotted. (F) Level of HRS and CD63 detected by western blot in EVs isolated from HeLa cells expressing either control or RL let-7a reporters. (G) Stability of miR-122 measured in Exosomes (EVs) and at Cellular level. HeLa cells which do not express miR-122 were transfected with miRNA-122 mimic and EVs (Exosome) were isolated 24h post transfection. Isolated EVs were incubated in a cell free medium for another 24h at 37°C. At regular intervals intact miRNA content were measured. Similarly cellular content of miR-122 were also measured. Values were compared against the amount of miRNA present at the beginning and plotted as percent remained. ns: non-significant, * P<0.05, **P<0.01, ***P<0.0001.P values were determined by paired t test.

These further corroborate the data obtained with RNA isolated from total cellular, Exosome or EV associated and microvesicle associated fractions. The increase in used miRNA let-7a was only found with EV-associated fractions in presence of its target RL-3xBulge ‒let-7a (Figure 2E). Control miR-16 did not show similar changes in its EV content in presence of let-7a reporter mRNA. As expected, we did not find any significant change in total EV content (as evidenced from GP63 and HRS content of EVs isolated from control and let-7a reporter mRNA expressing cells) (Figure 2F). We measured the stability of miRNA in cellular and EV associated pool. We transfect HeLa cells with exogenous mature miR-122 and residual levels of miR-122 present in cellular and EV-associated faction were compared. A higher turnover of miR-122 in EV was noted compared to the cellular miR-122 (Figure 2G). This data suggest effectiveness of EV-mediated miRNA export as a key mechanism of miRNA turnover in human cells as upon export the miRNAs become less stable in EVs and therefore effectively can clear the used miRNA from donor cell miRNA pool.

### Substrate mediated export of miRNA is ESCRT independent

MVBs play an important role in restricting miRNP recycling and activity [16, 17]. These vesicles can fuse with cell membranes to form exosomes in animal cells [33]. Maturation of early endosomes to MVBs is dependent on ESCRT, a multiprotein complex associated with MVBs. Depletion of ALIX or HRS proteins, the ESCRT components, had no considerable effect on EV-associated miRNA levels. siRNA mediated knockdown of SMPD2 increased cellular miRNA levels collectively in MDA-MB-231 cells. RNAi for ESCRT components and SMPD2 revealed that the substrate dependent exosomal export of miRNA is ESCRT independent but requires ceramide biosynthesis (Figure 3A-B).

**Figure 3.**
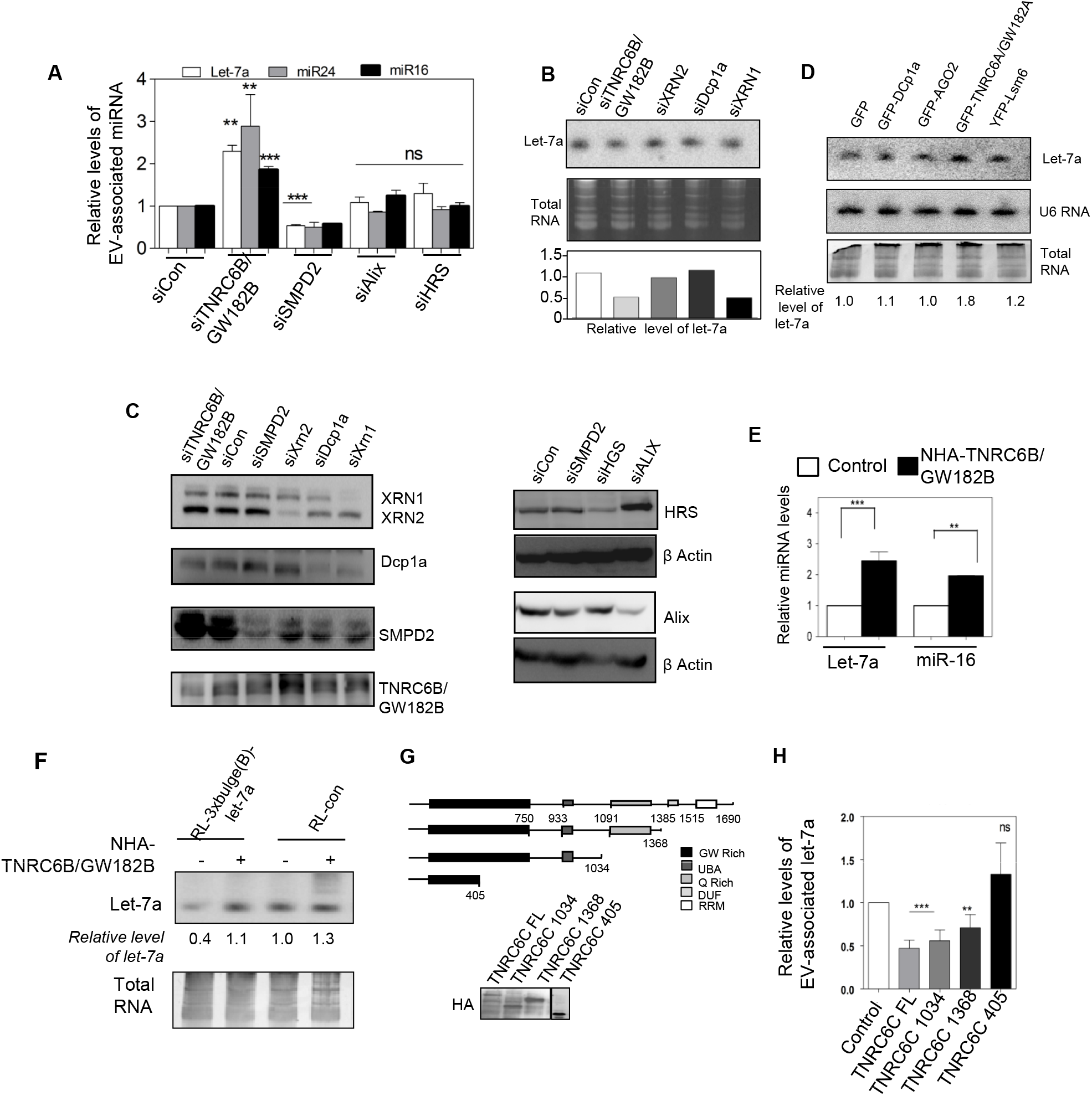
TNRC6/GW182 proteins restricts EV-mediated export of miRNA. (A) EV-mediated export of miRNA is ESCRT independent but require TNRC6B/GW182B. Realtime estimation of miRNA content in EVs from MDAM-MB-231 cells depleted for ESCRT components (ALIX and HRS), SMPD2 or TNRC6B/GW182B by RNAi (mean+/− s.e.m., n=3). (B) HeLa cells were transfected with siRNAs against different P-body components and levels of let-7a were determined by Northern blot. Relative level of let-7a in cells transfected with siRNAs were measured by real time quantification and mentioned below each lane. siRNA against RL luciferase was used as control. EtBr stained RNA gel used for blotting served as RNA loading control. Relative levels of let-7a are shown below (C) Western blotting for different P-body components was done with extracts prepared from respective siRNA treated cells to confirm effective knockdown. Levels of SMPD2 and ESCRT components ALIX and HRS in cells depleted for these factors by RNAi were detected. (D) Expression of P-body components alters let-7a level in HeLa cells. GFP-fusion constructs of different P-body components were transfected individually and level of let-7a in cells expressing these constructs were measured by Northern blot analysis. Total RNA was used as loading control. Relative levels of let-7a present in each condition are mentioned below the lanes. (E) Effect of NHA-TNRC6B/GW182B expression on cellular levels of two miRNAs miR-16 and let-7a in HeLa cells. (F) TNRC6B/GW182B restricts target dependent lowering of miRNA in MDA-MB-231 cells Levels of cellular let-7a in presence and absence of target reporter mRNA in cells also expressing NHA-TNRC6B/GW182B. Total RNA has been used as loading control and relative values of let-7a present are shown below each lane. (G-H) Ago2 interacting motifs of TNRC6C/GW182C is required for restricting miRNA export in human cells. The schematic diagram and expression of the full length and truncated versions of NHA-GW182C/GW182C in HeLa cells (G). Effect of expression of TNRC6C/GW182C and its truncated mutants on EV-associated let-7a level in MDAM-MB-231 cells (mean+/− s.e.m., n=4). ns: non-significant, * P<0.05, **P<0.01, ***P<0.0001.P values were determined by paired t test.

### GW182 proteins prevent target mediated lowering of miRNA

Ago interacting TNRC6/GW182 proteins are the effectors of miRNP mediated gene repression that can accelerate the decay of Ago bound mRNAs [25, 34–36] and may act as stabilizer of miRNPs in animal cells [26]. RNAi against TNRC6B/GW182B enhanced EV-mediated export of miRNA (Figure 3A-C). Contrary to increased extracellular export of miRNAs in human cells depleted for TNRC6B/GW182B, its ectopic expression restricted the extracellular export and increased cellular miRNA level (Figure 3D-E). Target dependent miRNA-lowering also got restricted in cells expressing TNRC6B/GW182B (Figure 3F). Expression of TNRC6C/GW182C (another member of TNRC6 protein family) also increased cellular miRNA level (Figure 3G-H). The C-terminal motif of TNRC6C, necessary for miRNA-mediated gene repression, was dispensable for its inhibitory role on EV-mediated export but the middle Ago2 interacting motif was found to be important (Figure 3G-H). Depletion or exogenous expression of majority of other GFP-tagged P-body components did not have any prominent effect on cellular miRNA level in HeLa cells except GFP-GW182A (Figure 3B-D).

### GW182 proteins prevent exosomal export of miRNA by retaining the miRNPs with P-Body

To understand the mechanism by which the TNRC6/GW182 proteins restrict EV-mediated export of used miRNAs, we separated the subcellular organelles on an Optiprep™ gradient to identify the fractions with which miRNPs remained associated in cells expressing NHA-TNRC6B/GW182B. In animal cells, miRNAs primarily remain associated with MVBs [16, 17] and ER [37]. With NHA-GW182 expression we have documented increased association of both Ago2 and miRNAs with Endosome/MVB enriched fractions in MDA-MB-231 and HeLa cells (Figure S2).

Relative subcellular distribution of miRNPs may get altered with changes in cell physiology and on availability of excess substrate RNA. Depletion of TNRC6B/GW182B by siRNA reduced P-body localization of Ago2 (Figure 4A-B). With NHA-TNRC6B/GW182B expression, we found an increased association of miRNA let-7a with MVB associated pool (Figure 4C-D). TNRC6B/GW182B expression increased the size of Ago2 positive P-bodies that showed a retarded rate of exchange of Ago2 molecules localized in bodies with the cytosolic pool (Figure 4 E-F). Therefore P-body compartmentalization of miRNPs in human cells, regulated by TNRC6/GW182 proteins, may control extracellular export and hence stability of miRNA in mammalian cells. Additionally, in NHA-TNRC6B/GW182B expressing human cells, Ago2 bodies were found to be less dynamic with reduced mobility (Figure S3 and Movie 1 and 2).

**Figure 4.**
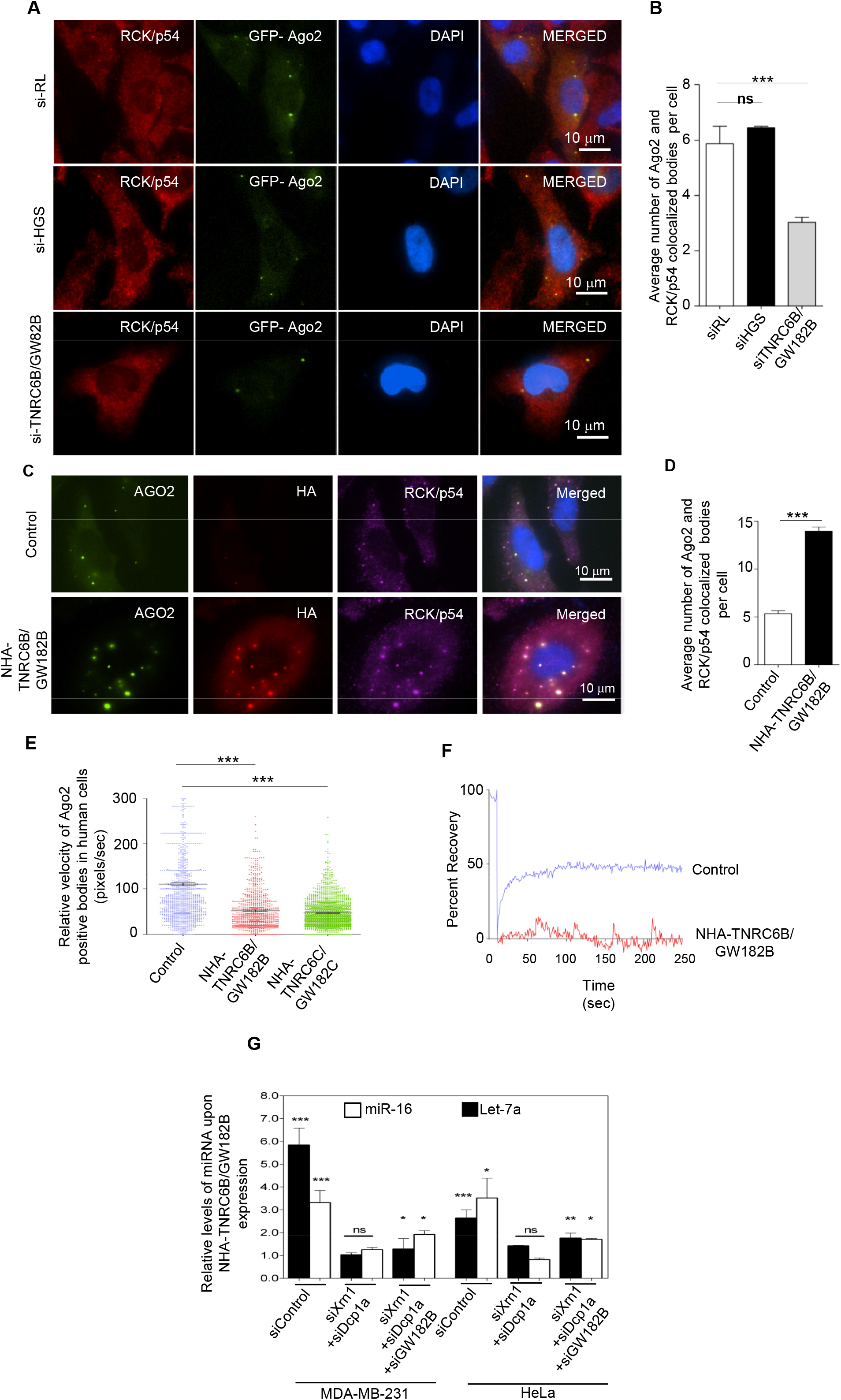
GW182 proteins promote P-body targeting of miRNPs to restrict the export. (A) Inhibition of TNRC6B/GW182B by RNAi reduces AGO2 positive bodies in HeLa cells. Cells were transfected with GFP-Ago2 and siRNAs either against TNRC6B/GW182B or HGS. GFP-AGO2 bodies were visualized in cells where they co-localized with RCK/p54. siRL was used as control (mean +/− s.e.m., n=20). (B) Quantification of Ago2 bodies showing co-localization with P-body marker RCK/p54 under different context (mean +/− s.e.m., n=10 cells, P=0.0005). (C) Larger and higher number of P-bodies in NHA-TNRC6B/GW182B expressing cells. HeLa cells were cotransfected with GFP-Ago2 encoding plasmid along with either the control or the NHA-TNRC6B/GW182 encoding plasmids. GFP positive Ago2 bodies were visualized and their localization with P-body marker RCK/p54 was documented. Increased size of Ago2 bodies showing colocalization with TNRC6B/GW182 was visualized in cells expressing NHA-TNRC6B/GW182B. (D) Relative quantification of colocalization of Ago2 and Rck/p54 from the experiments described in panel C, was done and plotted (mean +/− s.e.m., n=23 cells, P<0.0001). (E) From live cell time lapse imaging, the instantaneous velocity of multiple GFP-Ago2 particles were measured and plotted for both control and NHA-TNRC6B/GW182B plasmids transfected cells (mean +/− s.e.m., n=600, P<0.0001). (F) Impaired recovery of GFP fluorescence of GFP-Ago2 in bodies after photobleaching in cells expressing NHA-TNRC6B/GW182B with respect to control. (G) Effect of depletion of PBs on miRNA levels in cells expressing NHA-TNRC6B/GW182B. The protein was expresed in HeLa cells prior-targetted with siRNAs against different P-body components. The cellular miRNA levels in individual cases were measured against control cells tranafected with control siRNA (siRL) (mean+/− s.e.m., n=4). ns: non-significant, * P<0.05, **P<0.01, ***P<0.0001.P values were determined by paired t test.

The dependence of miRANA stabilization by GW182 found to be dependent on integrity of PBs, and depletion of PBs by siRNAs against specific components like Xrn1 and Dcp1a resulted in reduced stabilization of miRNA in HeLa and MDA-MB-231 cells expressing NHA-TNRC6B/GW182B (Figure 4G).

In this context HuR is known to delocalize miRNA repressed messages from PBs and ensure its translation in stressed human cells without an effect on PBs [23]. HuR is also know to facilitate the export of miRNAs under stress condition or when expressed ectopically in human cells by reversibly binding the miRNA in hepatic cells and ubiquitination of HuR is the key step happening on endosomes that allows its miRNA unbinding and loading of miRNA in MVBs for EV-mediated export [21]. Could GW182 protein counteract the miRNA export mediated by HuR? GW182 found to facilitate the relocalization of miRNPs to PBs and this phase separated miRNPs are unavailable for export. In this process of target RNA driven export, the interaction of GW182 with Ago2 happens primarily on the endsomal fraction [24] and therefore a event most likely happen upon relocalization of the repressed mRNAs along with miRNPS to the endosome. HuR on contrary work and ER attached polysome first to disengage the miRNPs from the targets and then to uncouple miRNA from Ago2 [21]. Therefore these two processes are likely to be independent to each other. We have observed a limited effect of HuR over expression on P-body size and Number (Figure 5 A-C) and has little effect on Ago2 mobility in mammalian cells (Figure 5D). HuR over expression accelerate miRNA export and siRNA mediated GW182 depletion or over expression of NHA-TNRC6B/GW182B don’t seems to have any significant effect on the cellular miRNA content already lowered in HuR expressing cells. These results confirmed the uncoupling of GW182 mediated miRNA stabilization step from HuR mediated export of miRNA. Therefore, in mammalian cells, HuR mediated miRNA export is a predetermined step that can’t be reversed by Ago2 interacting proteins GW182 that work downstream of miRNP coupling of target message and happens in a spatio-temporally distinct manner than HuR driven miRNA export which is a target RNA independent event (Figure 5G).

**Figure 5.**
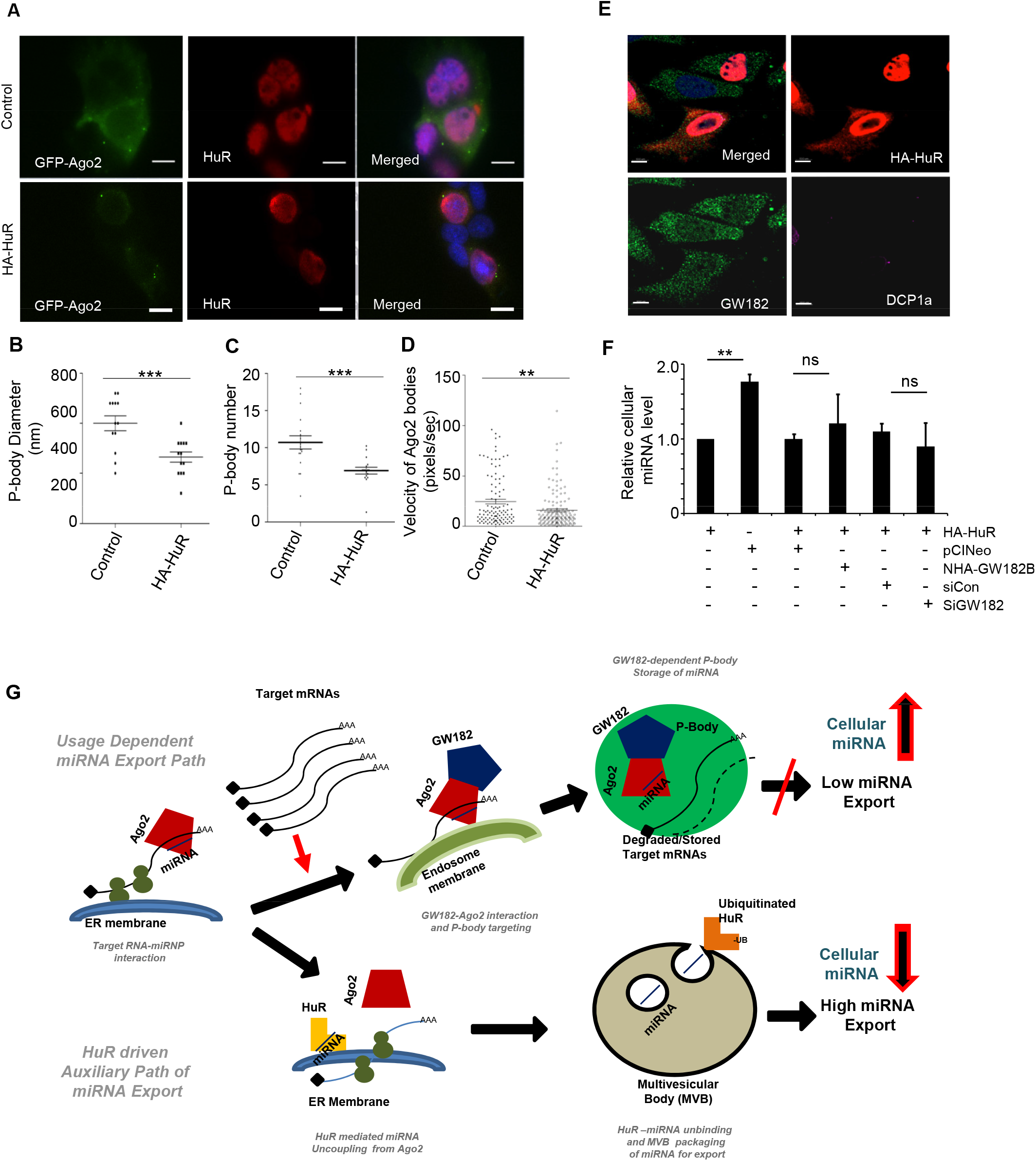
HuR mediated miRNA export is insensitive to cellular GW182 levels. **(A)** Effect of HA-HuR on Ago2-positive bodies in Huh7 cells. P-body localization of GFP-Ago2 in HA-HuR expressing cells. Cells were transfected with GFP-AGO2 encoding plasmid along with either the HA-HuR expression plasmid or control vector and GFP-positive bodies were visualized. Anti HuR antibody was used to detect HuR and DAPI was used to stain nucleus. (B-D) A scatter plot depicting HA-HuR mediated changes in diameter (mean +/− s.e.m.,n=18, P=0.0007) (B), number (mean +/− s.e.m., n=14, P=0.0004) (C) and mobility of GFP-AGO2 positive bodies in Huh7 cells (mean +/− s.e.m., n=108, P=0.0016) (D). (E) Effect of HA-HuR expression (Red) on Dcp1a (Purple) and GW182 (Green) in HeLa cells. (F) Effect of GW182 depletion or NHA-TNRC6B/GW82B expression on cellular miRNA export by HuR in HeLa cells. Levels of cellular miRNA in pCIneo or HA-HuR expressing HeLa cells were compared. In HA-HuR expression context, siGW182B treatment or NHA-GW182B expression seems to have no effect on cellular miRNA levels compared to control cells treated with siCon or expressing equivalent amount of pCIneo control plasmid (mean+/− s.e.m., n=4). ns: non-significant, * P<0.05, **P<0.01, ***P<0.0001.P values were determined by paired t test. Scale Bar 10 μm.

It has been shown before that blockage of EV-mediated export of miRNA-let-7a gets impaired in HuR compromised condition or in growth retarded human cells. The increase of let-7a induces cell senescence in human breast cancer cells MDA-MB-231 [21]. Does GW182 expression can also increase the senescence in MDA-MB-231 by enhancing cellular content of let-7a? As expected, we have noted enhanced cell proliferation and decrease in senescence in MDA-MB-231 cells depleted for GW182B supporting a role of GW182 protein in cell growth regulation (Figure S4).

## Discussion

miRNAs are otherwise considered as stable molecules with slow turnover rates in mammalian cells. Our findings provide evidence that miRNAs engaged in translational repression could be selected for export in target RNA dependent manner in human cells. Here, we focused on how the usage related regulated export of miRNA controls gene expression in mammalian cells. Our data also provides an important molecular link between usage of a miRNA and its extracellular export that can explain why export and hence turnover of miRNAs are selective in animal cells. In this work we have been able to document EV-mediated miRNA export as a mechanism used by human cells to regulate cellular miRNA content and have delineated the molecular mechanism underlying this phenomenon (Figure 5G).

In the presence of its substrates, miRNA is more susceptible to extracellular export and their levels are reduced in human cells. Previous observation in *C. elegans* suggests that the let-7 miRNA stability is dependent on its binding to the substrate [38]. This rationale may be valid for certain exceptional cases, but it is more common that target mRNAs and miRNAs are usually present in animal cells either in a mutually exclusive manner or their expressions are inversely correlated [39]. Furthermore, miRNAs have been reported to be “sponged” with mRNAs having multiple miRNA sites and are used to titrate out endogenous miRNAs to rescue their target genes accompanied with reduction of respective miRNAs [40]. Competing endogenous RNAs (ceRNAs) act as natural miRNA sponges and can attenuate the function of specific miRNAs by binding in a competitive manner to rescue the expression of the endogenous target genes in animals and plants [41, 42]. It is also interesting to test whether ceRNAs can accelerate the export of miRNAs in animal cells. Besides non-coding RNAs, developmental stages of lymphopoiesis, has also been reported to modulate miRNA abundance brought about by changes in mRNA profiles during ontogeny [14]. Modification of miRNA with additional bases in its 3’end is also known to play important role to modulate miRNA stability and activity in animal cells [43, 44]. Whether such a modification in a substrate dependent manner can modulate miRNA export will be a subject of interest for future exploration.

In this context the data published by Squadrito et al. [45] suggests an apparent opposite effect of targets on the miRNAs having the binding sites. They have argued in favour of retention of miRNAs in PBs in presence of the target mRNAs and thus exclusion of respective miRNAs from exosome associated miRNA pool. This could be a cell type specific event and considering an abundant amount of GW182 may present in the macrophage system, miRNP storage in PBs could happen in a target dependent manner but should not hold true for the proliferative cancer cells where rapid turnover of transcripts are also ensured (Figure 5G).

Alternative to extracellular degradation of exosomal miRNAs, miRNA-loaded EVs are used for intercellular communications where the retaken EV contents are reused in recipient cells [18, 30]. The EV release of used miRNAs may ensure the homogeneity of miRNA expression and activities among neighbouring cells. In a tissue, the spiked expression of target mRNAs in individual cells in a population may be tamed by the extra miRNAs received via EVs till the balanced expression of target genes and miRNAs are achieved. This possibly ensures robustness of the biological processes in a tissue as was discussed by Ebert and Sharp elsewhere [46].

We have identified TNRC6/GW182 and HuR as inverse regulators of the exosomal export process and it would be interesting to identify additional factors that may act as regulators of miRNA export. miRNA homeostasis is controlled by several factors that contribute in buffering of cellular miRNA level and stability in animal cells. From this study it also seems that P-bodies can effectively act as a reservoir for used miRNAs that otherwise get exported or recycled via MVB/exosomes (Figure 5G). Increase of let-7a in breast cancer cells impaired for EV-mediated export increased senescence and sensitized cells to anti cancer drug. Therefore, the manipulation of miRNA turnover machineries may lead to a better management of anti proliferative drugs in cancer treatment.

## Experimental Procedures

### Expression plasmids, cell culture and transfection

Information of all plasmids and siRNAs used in this study are available in the Table S1. Information of all DNA oligos used as primers for real time quantification and gene amplification are listed in Table S2. All human cell lines were grown in Dulbecco’s modified Eagle’s medium (DMEM) supplemented with 2 mM L-glutamine and 10% heat-inactivated fetal calf serum (FCS). For starvation experiments, cells were incubated in Hanks’ Balanced Salt Solution (HBSS) supplemented with 10% dialyzed FCS (all from Invitrogen). SMPD2 inhibitor GW4869 was used at a final concentration of 10 μM (Calbiochem). For all experiments cells were grown to 25-40% confluency states unless specified otherwise.

Transfections of HeLa, MDA-MB-231 and Huh7 cells were all were performed using Lipofectamine 2000 (Invitrogen) according to manufacturer’s protocol. For miRNA repression assays 25 ng Renilla Luciferase (RL) reporter plasmids with 500 ng Firefly Luciferase (FL) plasmid were introduced per well of a six-well cell culture format (10 cm^2^ area). Luciferase activities were measured after 48h of transfection with a cell splitting after 24h post-transfection. For target RNA or HA-HuR overexpression experiments, 2 μg of respective plasmids were transfected per well of a six well plate and done in triplicate. Inducible expression of let-7a and miR-122 reporter plasmid was achieved by incubation with 300 nm of Tetracycline in HeLa cells grown in Tetracycline free FCS and also expressing a Tet-repressor protein from the pTet-On Advance plasmid introduced before. siRNAs were transfected at 50 nM concentration using either Lipofectamine RNAiMAX or Lipofectamine 2000. AllStars Negative Control siRNA (Qiagen) or a siRNA against Renilla Luciferase (siRL) were used as control (siCon).

### Luciferase assay, Northern and Western blot

Renilla Luciferase (RL) and Firefly Luciferase (FL) activities were measured using a Dual-Luciferase Assay Kit (Promega) following the suppliers protocol on a VICTOR X3 Plate Reader with injectors (Perkin Elmer). FL normalized RL expression levels for reporter and control were used to calculate fold repression as the ratio of control to reporter normalized RL values.

Northern blotting of total cellular RNA (5-15 μg) was performed as described earlier [23, 28]. For miRNA detection, ^32^P-labeled 22 nt anti-sense DNA or LNA modified probes specific for respective miRNAs or siRL or U6 snRNA were used. PhosphorImaging of the blots was performed in Cyclone Plus Storage Phosphor System (Perkin Elmer) and Optiquant software (Perkin Elmer) was used for quantification.

Western analyses of different miRNP components were performed as described previously (20). Detailed list of antibodies used for western blot are available in Table S3. Imaging of all western blots was performed using an UVP BioImager 600 system equipped with VisionWorks Life Science software (UVP) V6.80.

### Preparation of Extracellular Vesicles (EV) and treatments

EVs were isolated based on the published protocols [21, 47]. Briefly, for isolation of EVs, cells were grown in medium precleared for the same and the culture supernatants were clarified for cellular debris and other contaminants by centrifugation at 400xg for 5 min, 2,000xg for 10 min, and 10,000xg for 30 min and filtration through a 0.22 μm filter. EVs designated as exosomes were separated by centrifugation at 100,000xg for 75 min and were resuspended in 200μl of 1X passive lysis buffer (Promega) for isolation of exosomal proteins and RNA. Characterization of isolated exosome and its purity check were done as the procedures adopted elsewhere [21].

### Fractionation and separation of Endosomes and ER on Optiprep gradients

Optiprep™ (Sigma-Aldrich, USA) was used to prepare a 3-30% continuous gradient in a buffer containing 78 mM KCl,4 mM MgCl_2_,8.4 mM CaCl_2_,10 mM EGTA,50 mM HEPES-NaOH (pH 7.0) for separation of subcellular organelles as described previously with minor modifications described below [48]. Cells were trypsinized, washed and homogenized with a Dounce homogenizer in a buffer containing 0.25 M sucrose,78 mM KCl,4 mM MgCl_2_,8.4 mM CaCl_2_,10 mM EGTA,50 mM HEPES-NaOH pH 7.0 supplemented with 100 μg/ml of Cycloheximide 5 mM Vanadyl Ribonucleoside Complex (VRC) (Sigma Aldrich), 0.5 mM DTT and 1X Protease Inhibitor. The lysate was clarified by centrifugation at 1,000xg for 5 minutes and layered on top of the prepared gradient. The tubes were centrifuged for separation of gradient using established protocols and ten fractions were collected.

### Immunofluorescence

For immunofluorescence, cells were transfected with 250 ng of GFP-Ago2 alone or co-transfected with 2 μg of either NHA-TNRC6B/GW182B (or HA-HuR) expression plasmid per well of a six well plate. pCIneo plasmid was used as a control. The cells were split after 24h of transfection and subjected to specific experimental conditions. For immunofluorescence analysis, cells were fixed with 4% paraformaldehyde for 30 min, permeabilized and blocked with PBS containing 1% BSA and 0.1% Triton X-100 and 10% goat sera (GIBCO) for 30 min. Secondary anti-rabbit or anti-mouse antibodies labeled either with Alexa Fluor® 488 dye (green), Alexa Fluor® 594 dye (red) or Alexa Fluor® 647 dye (far red) fluorochromes (Molecular Probes) were used at 1:500 dilutions. Cells were observed under a Plan Apo VC 60X/1.40 oil or Plan Fluor 10X/0.30 objectives on an inverted Eclipse Ti Nikon microscope equipped with a Nikon Qi1MC or QImaging-Rolera EMC^2^ camera for image capture.

For live cell analysis 250 ng of GFP-Ago2 encoding plasmid was used for transfection of a six well plate which was further transfected with NHA-GW182 or HA HuR after overnight durations. pCIneo was used as the control plasmid. The cells were split at 24h post transfection period for analysis. For FRAP experiments, cells were photobleached, and recovery was monitored for a total duration of less than 4 minutes. Glass bottom petridishes pre-coated with gelatin were used for cell growth for live cell imaging. Cells were observed with a 60X/N.A.1.42 Plan Apo N objective. Images were captured with a IXON3 EMCCD camera in an ANDOR Spinning Disc Confocal Imaging System on an Olympus IX81 inverted microscope.

### Analyses of mRNA and miRNA by qRT-PCR

Real time analyses by two-step RT-PCR was performed for quantification of miRNA and mRNA levels on a 7500 REAL TIME PCR SYSTEM (Applied Biosystems) or Bio-Rad CFX96™ real time system using Applied Biosystems Taqman chemistry based miRNA assay system. mRNA real time quantification was generally performed in a two step format using Eurogentec Reverse Transcriptase Core Kit and MESA GREEN qPCR Master Mix Plus for SYBR Assay with Low Rox kit from Eurogentec following the suppliers’ protocols. The comparative C_t_ method which typically included normalization by the 18S rRNA levels for each sample was used for relative quantification.

miRNA assays by real time PCR was performed with 25 ng of cellular RNA and 200 ng of exosomal RNA unless specified otherwise, using specific primers for human let-7a (assay ID 000377), human miR-122 (assay ID 000445) human miR-16 (assay ID 000391). U6 snRNA (assay ID 001973) was used as an endogenous control. One third of the reverse transcription mix was subjected to PCR amplification with TaqMan® Universal PCR Master Mix No AmpErase (Applied Biosystems) and the respective TaqMan® reagents for target miRNA. Samples were analyzed in triplicates from minimum two biological replicates. The miRNA levels were defined from the cycle threshold values (C_t_) for representation of exosomal miRNA or immunoprecipitated miRNA levels. The comparative C_t_ method which typically included normalization, by the U6 snRNA, or a non relevant miRNA, for each sample was used for all other instances.

### Cell senescence and TUNEL assay

Cells were assayed for senescence using Senescence Cells Histochemical Staining Kit from Sigma after fixation of the cells in a buffer containing 20% formaldehyde, 2% glutaraldehyde, 70.4 mM Na_2_HPO_4_, 14.7 mM KH_2_PO_4_, 1.37 M NaCl, and 26.8 mM KCl as a stock 10% solution, for 7 mins. TUNEL assays were performed with DeadEnd™ Fluorometric TUNEL System kit (Promega) as per manufacturer’s protocol. Cells were mounted on a cover slide with Vectashield DAPI for observation with a 10X Plan Fluor 10X/0.30 objective on a Nikon Eclipse Ti microscope and imaged on Nikon Qi1MC camera.

### Post imaging analysis & others

All western blot and Northern blot images were processed with Adobe Photoshop CS4 for all linear adjustments and cropping. All images captured on Nikon Eclipse Ti microscope or ANDOR spinning disc microscope were analyzed and processed with Nikon NIS ELEMENT AR 3.1 software including P-Body tracking and velocity calculations. Image cropping was done using Adobe Photoshop CS4. All antibodies used in this study are listed in Supplementary Table 3. Statistical analysis of data were done by performing unpaired, non-parametric, two-tailed t-tests with a confidence interval of 95% and four significant digits were considered in all cases.

## Author Contributions

The work is conceived and analyzed by S.N.B and he was helped by other authors in formulating, analyzing and critical reading of the text and Figures. S.G., Y.C., K.M. and B.G did all the experiments and data analysis. K.M. and S.G. also contributed in conceptualization of the idea.

## Acknowledgements

We would like to thank W Filipowicz, G Meister, R Pillai, E Bertrand for their generous help with reagents and plasmids constructs. We also like to thank B. Barman who helped us with plasmid construction. The work has been primarily funded by Swarnajayanti Fellowship and CEFIPRA grant to S.N.B. All the authors except S.N.B and K.M. were supported by fellowship from CSIR.

## Supplementary Documents

### Supplementary Figure Legends

**Figure S1.**
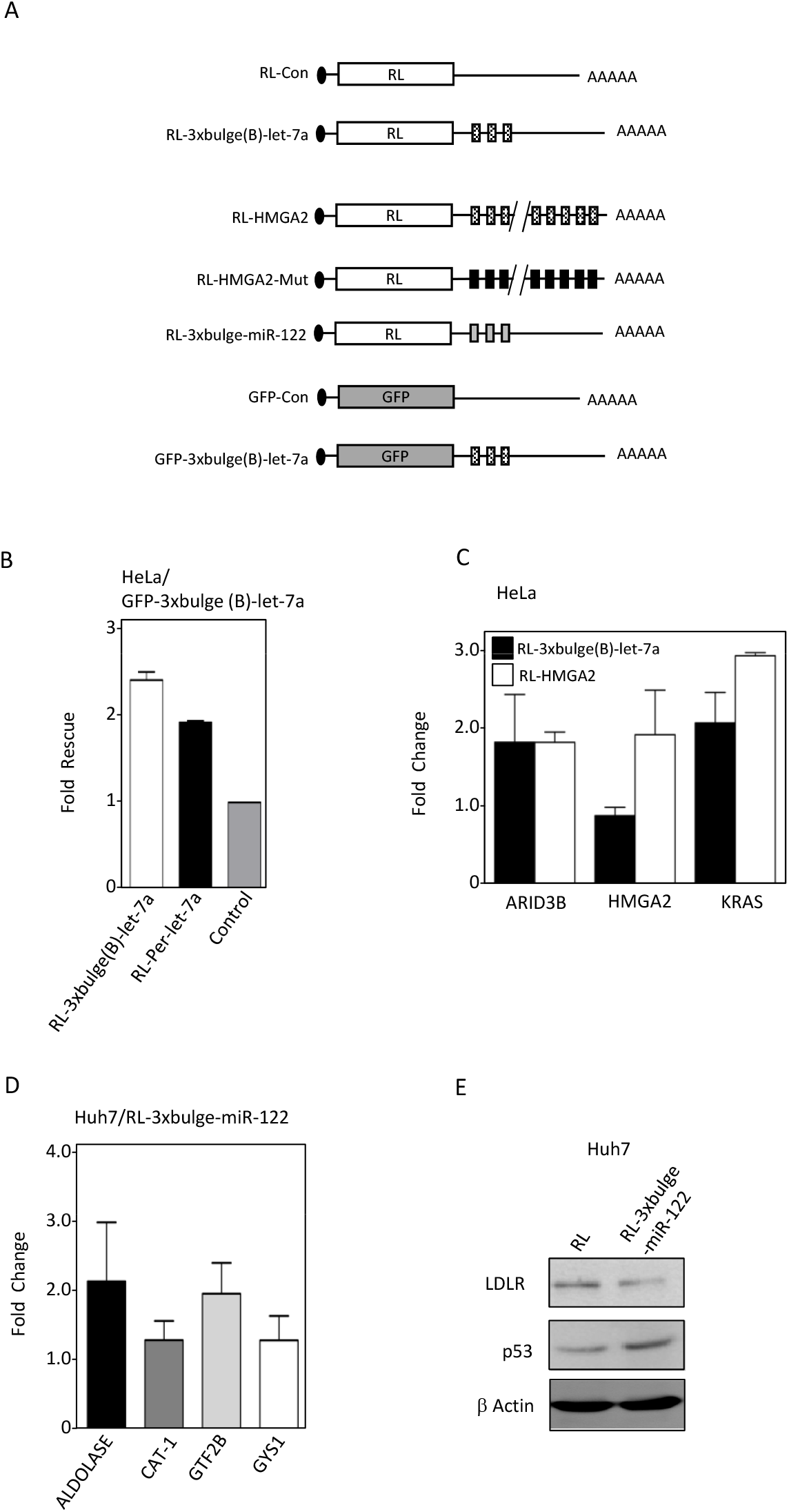
Reversal of miRNAs action in mammalian cells expressing excess target mRNAs, related to Figure 1. (A) Scheme of target mRNAs used for reversal of miRNA action endogenous targets. (B) Derepression of let-7a reporters in HeLa cells. Normalized expression level of different RL reporters were measured in cells coexpressing GFP mRNAs with or without three let-7a target sites. Fold rescue represents the fold increase in activity of each RL reporter estimated by RL protein expressed in GFP-let-7a expressing cells than GFP-con. (C-D) Expression of target mRNA leads to upregulation of endogenous miRNA targets. Fold increase of endogenous target mRNAs in HeLa (C) and Huh7 (D) cells expressing let-7 (RL-3xbulgeB-let7a or RL-HMGA2) and miR-122 (RL-3xbulge-miR-122) reporter were estimated by real time quantification against their expression levels in control reporter transfected cells. For RL-HMGA2, the control was RL-HMGA2-Mut reporter with mutated let-7a binding sites, while RL-con served as control for the rest. (E) Reduction in miRNA level by its substrate leads to derepression of endogenous targets at protein level. Expression of a reciprocally regulated indirect miR-122 target, LDL Receptor and the direct target p53 in Huh7 cells expressing reporters with or without miR-122 sites. Β-Actin serves as loading control.

**Figure S2.**
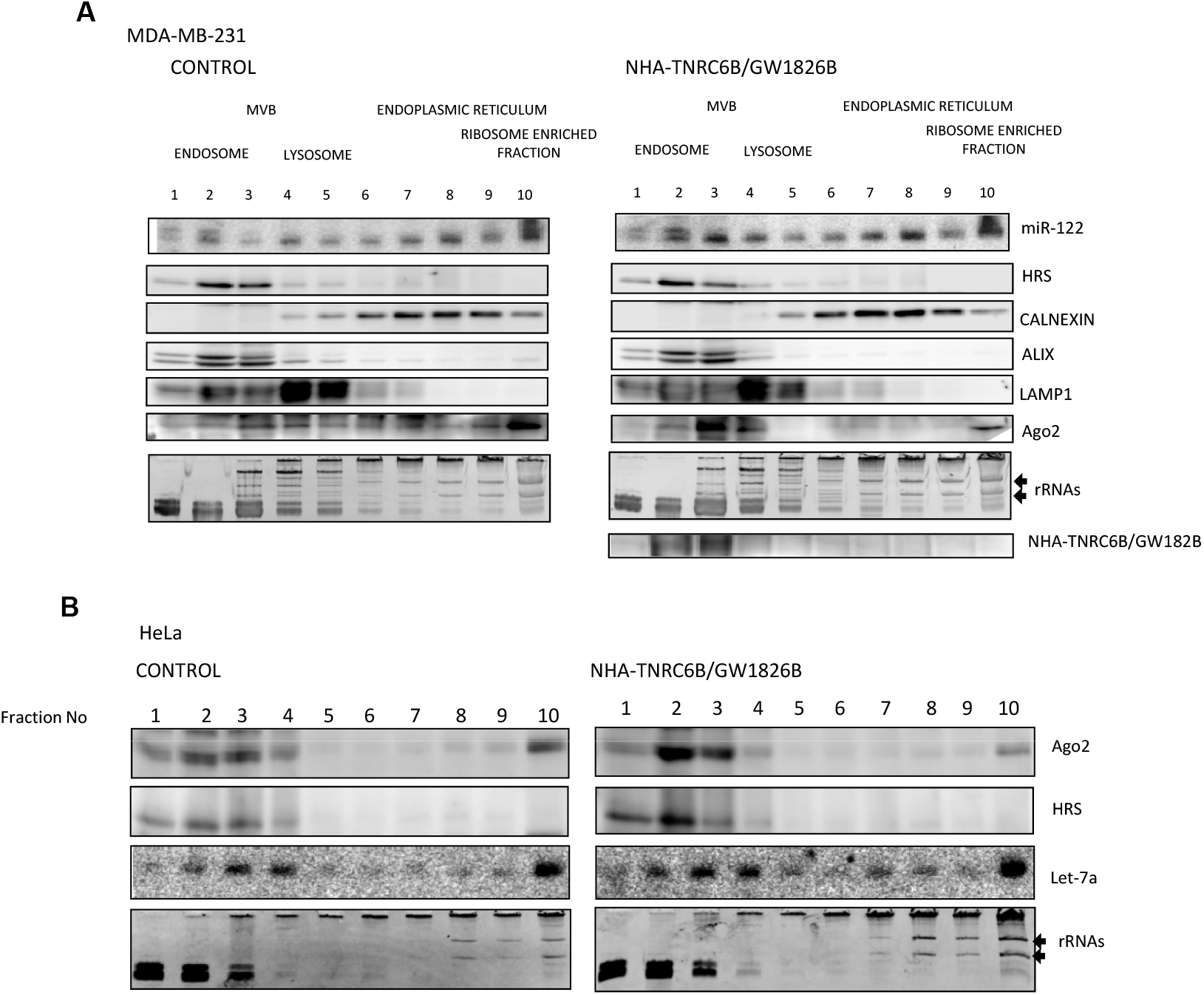
Distribution of Ago2 and miRNA in MDA-MB-231 and HeLa cells expressing TNRC6B/GW182B. Increased miRNP is associated with endosome faction positive for HRS marker protein in NHA-TNRC6B/GW182B expressing MDA-MB-231 (A) and HeLa (B) cells. The 3-30% Optiprep™ gradient analysis and distribution of Ago2 and in cells expressing NHA-TNRC6B/GW182B. Expression and distribution of NHA-TNRC6B/GW182B was determined by Western blot using anti-HA antibody in panel A. Levels of let-7a or exogenously expressed miR-122 has been determined by Northern Blot analysis. Calnexin serve as ER marker. Alix is also an endosomal marker while Lamp1 is a lysozomal marker.

**Figure S3.**
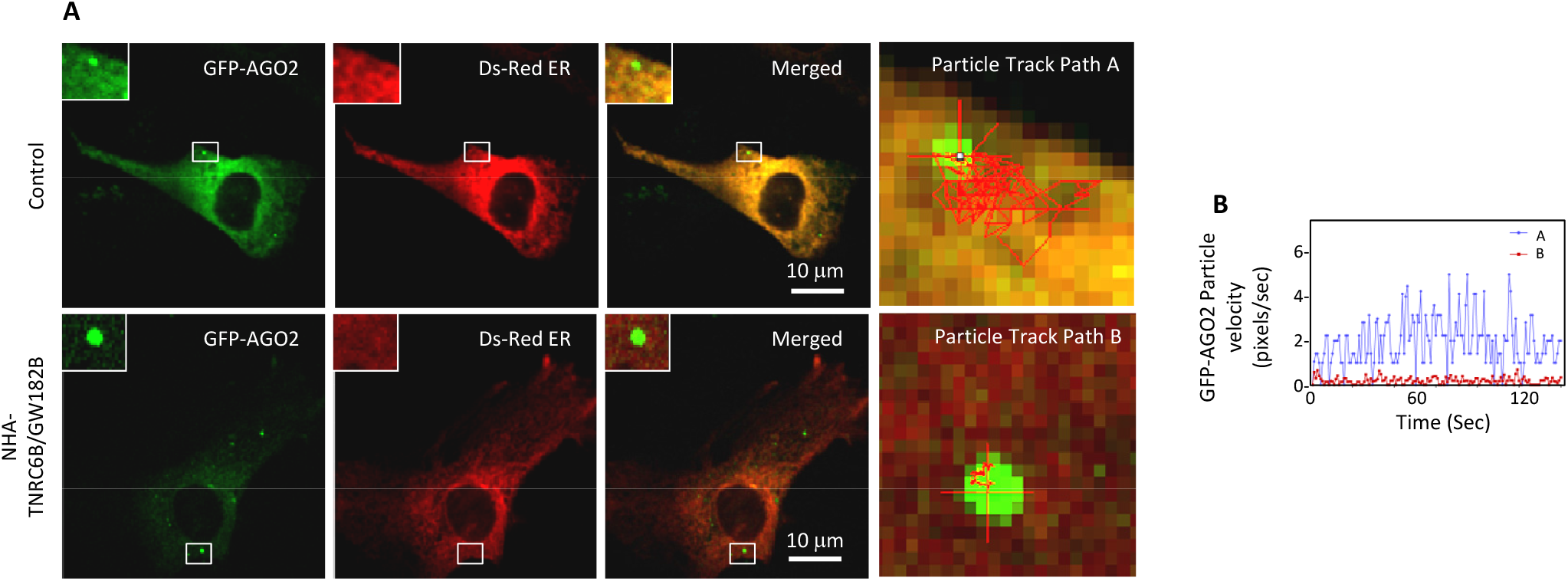
Lower mobility of Ago2 bodies in NHA-TNRC6B/GW182B expressing cells. **(A)** Localization of GFP-AGO2 bodies in cells coexpressing NHA-TNRC6B and or control vector. **(B)** Lower mobility of Ago2 bodies. Change in particle velocity with time for two GFP-AGO2 positive bodies in control (Path A, Panel A) and NHA-TNRC6B/GW182B expressing (Path B, Panel A) HeLa cells.

**Figure S4.**
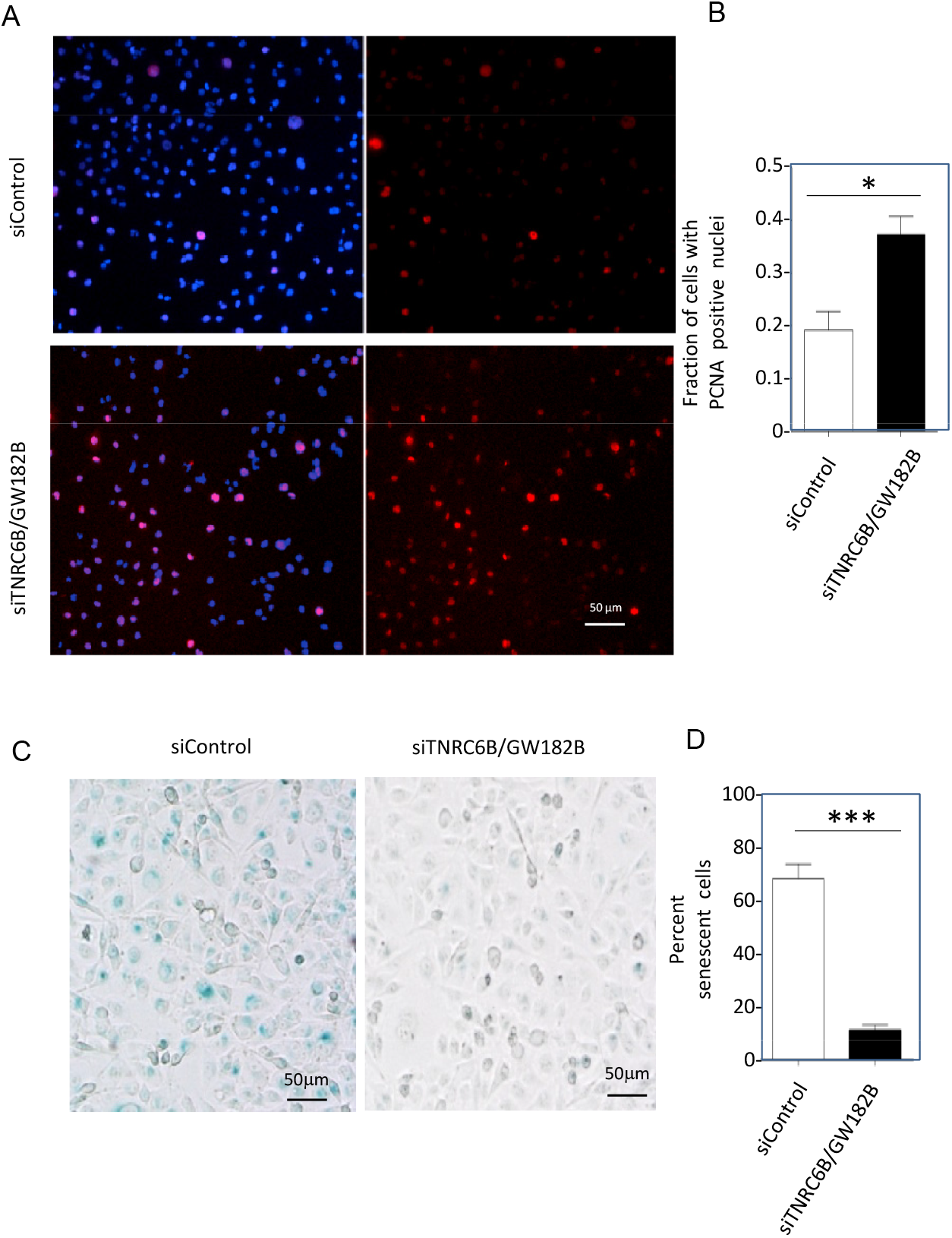
Increased senescence and decreased proliferation of MDA-MB-231 cells treated with siGW182B. (A-D) Increased number of proliferating cells with PCNA positive nuclei in siTNRC6B/GW182B treated cells was visualized (A) and plotted (mean +/− s.e.m., n=2 independent experiments, P=0.0212) (B).Reduced senescence in cells treated with siTNRC6B was detected (C) and quantified (mean +/− s.e.m., n=2 independent experiments, P=0.007) (D).

**Supplemental Movies S1 and S2 Time Lapse imaging of GFP-AGO2 in HeLa cells expressing dsRED-ER, transfected with control (Movie S1) or NHA-GW182B (Movie S2).** Cells were subjected to time lapse imaging analysis for 2 minutes each with a total of one hundred and fifty images being captured at equally spaced time intervals. Frame Rate: 4 frames per second.

### Supplemental Tables

**Table S1.**
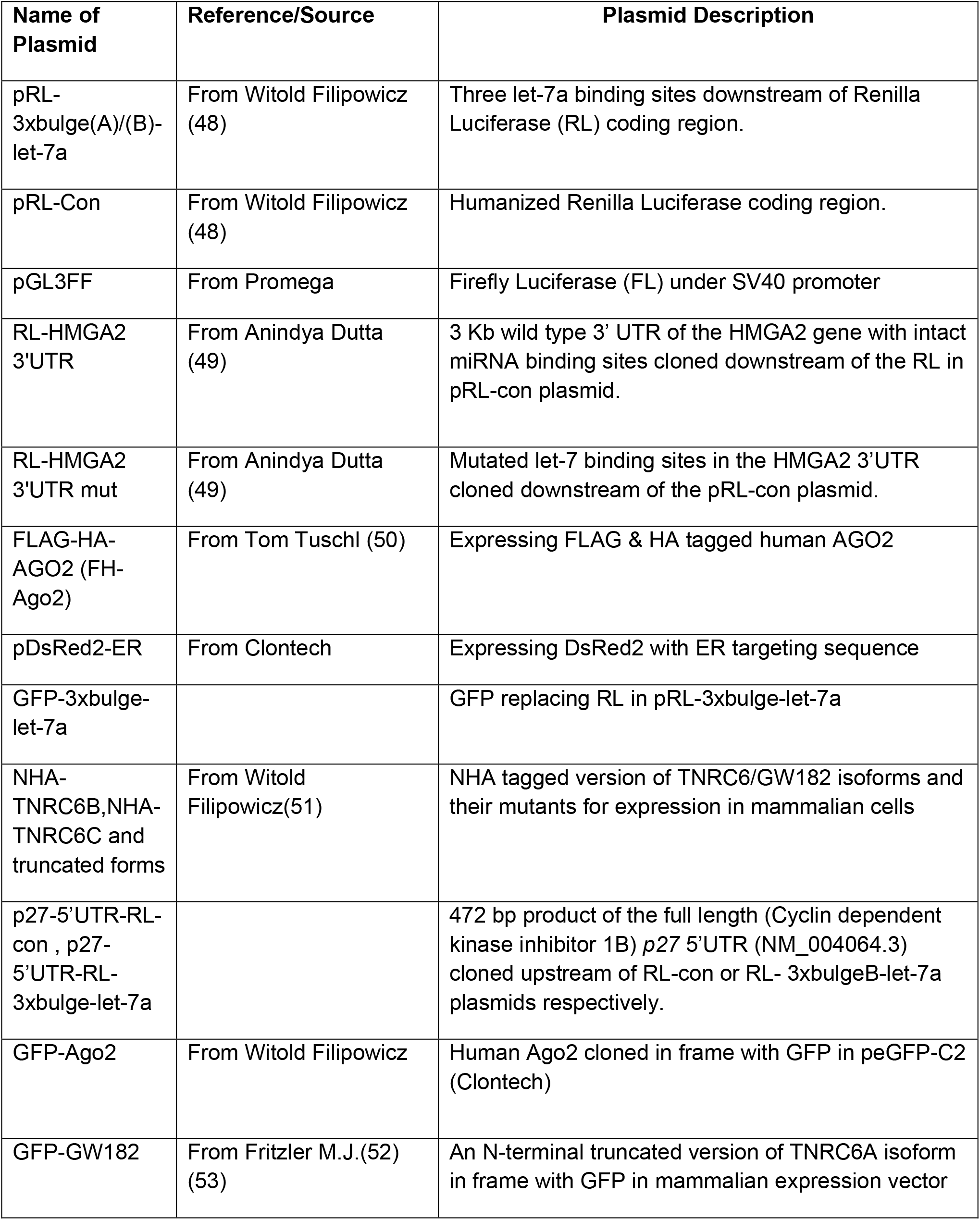
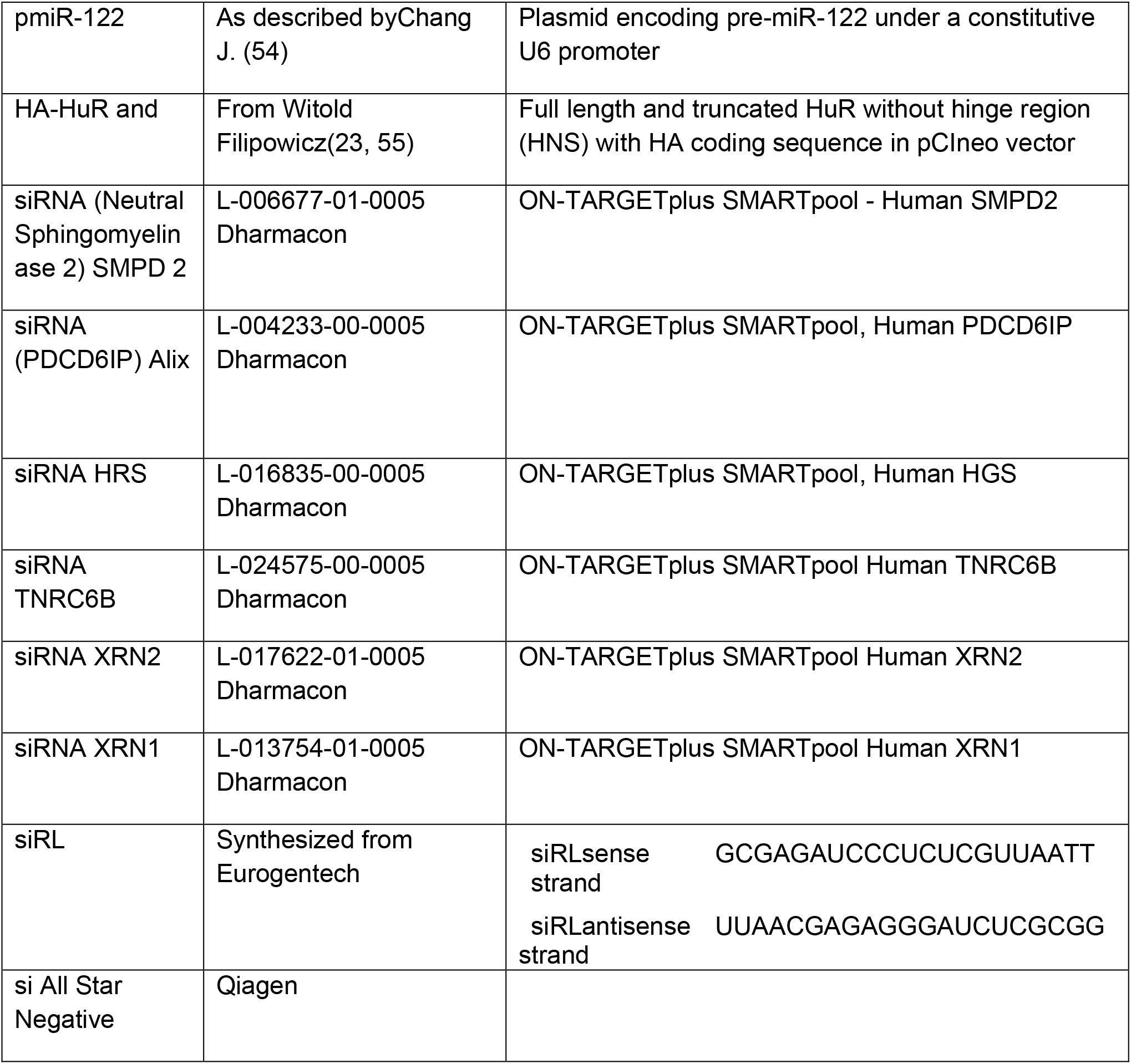
List of Plasmids and siRNA.

**Table S2.**
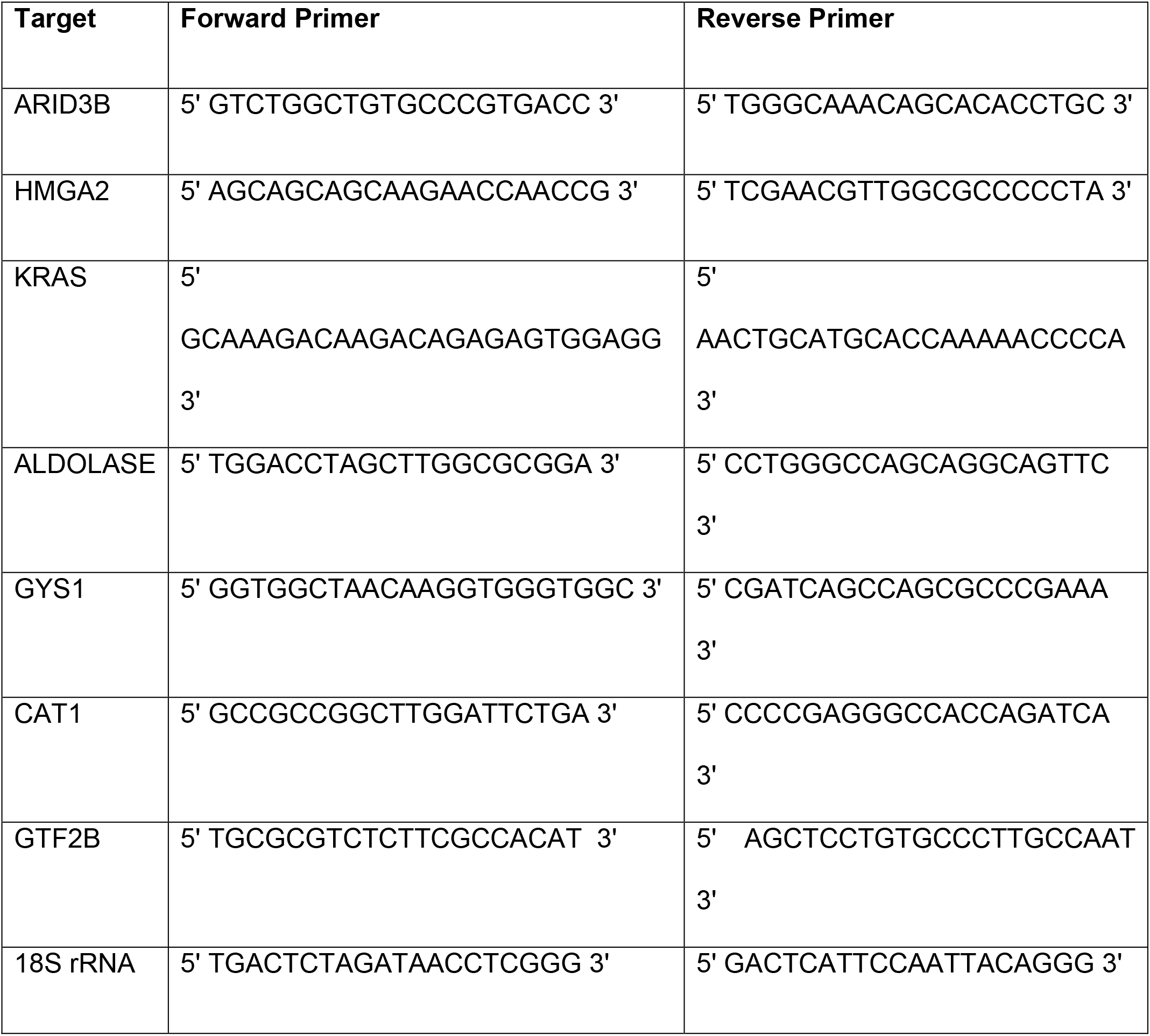
List of primers used.

**Table S3.** List of Antibodies used.

| **Name of Antigens** | **Raised in** | **Source** | **Dilutions used for Western Blot** |
| --- | --- | --- | --- |
| p53 | Rabbit | Biovision | 1:1000 |
| LDL Receptor | Rabbit | Cayman Chemical | 1:1000 |
| PCNA | Rabbit | Bethyl Laboratories | 1:1000 |
| Hsp90 | Rabbit Monoclonal | Cell Signalling | 1:1000 |
| β-Actin | Mouse Monoclonal<br>(HRP conjugated) | Sigma Aldrich | 1:10000 |
| HA | Rat Monoclonal<br>( clone 3F10) | Roche | 1:1000 |
| GFP | Mouse Monoclonal<br>(clones 7.1 and 13.1) | Roche | 1:1000 |
| Ago2<br>(eIF2C2) | Mouse Monoclonal | Novus Biologicals | 1:500 |
| XRN1 | Rabbit | Bethyl Laboratories | 1:10000 |
| RCK/p54 | Rabbit | Bethyl Laboratories | 1:10000 |
| Dcp1a | Mouse Monoclonal | Novus Biologicals | 1:1000 |
| HRS | Rabbit | Bethyl Laboratories | 1:1000 |
| $\beta$ Tubulin | Mouse | Sigma | 1:1000 |
| Calnexin | Rabbit | Cell Signalling | 1:1000 |
| Alix | Mouse Monoclonal | Santa Cruz | 1:500 |
| GW182 | Rabbit | Bethyl Laboratories | 1:1000 |
| XRN2 | Rabbit | Bethyl Laboratories | 1:10000 |
| SMPD | Mouse Monoclonal | Abnova | 1:2000 |
| HuR | Mouse Monoclonal<br>(clone 3A2) | Santa Cruz | 1:1000 |

